# Targeted reconstruction of T cell receptor sequence from single cell RNA-sequencing links CDR3 length to T cell differentiation state

**DOI:** 10.1101/072744

**Authors:** Shaked Afik, Kathleen B. Yates, Kevin Bi, Samuel Darko, Jernej Godec, Ulrike Gerdemann, Leo Swadling, Daniel C. Douek, Paul Klenerman, Eleanor J. Barnes, Arlene H. Sharpe, W. Nicholas Haining, Nir Yosef

**Affiliations:** Computational Biology Graduate Group, UC Berkeley, Berkeley, CA, USA; Department of Pediatric Oncology, Dana-Farber Cancer Institute, Harvard Medical School, Boston, MA, USA; Human Immunology Section, Vaccine Research Center, NIAID, NIH, Bethesda, MD, USA; Department of Microbiology and Immunobiology, Harvard Medical School, Boston, MA, USA; Evergrande Center for Immunologic Diseases, Harvard Medical School and Brigham and Women’s Hospital, Boston, MA, USA; Translational Gastroenterology Unit, Peter Medawar Building for Pathogen Research, University of Oxford, Oxford, UK; NIHR Oxford Biomedical Research Centre, John Radcliffe Hospital, Oxford, UK; Broad Institute of MIT and Harvard, Cambridge, MA, USA; Division of Hematology/Oncology, Children’s Hospital, Harvard Medical School, Boston, MA, USA; Department of Electrical Engineering and Computer Science and Center for Computational Biology, UC Berkeley, Berkeley, CA, USA; Ragon Institute of Massachusetts General Hospital, MIT and Harvard, Cambridge, MA, USA

**Author notes:** These authors contributed equally (Shaked Afik, Kathleen B. Yates, Kevin Bi). These authors contributed equally (W. Nicholas Haining, Nir Yosef).

## Abstract

The T cell compartment must contain diversity in both TCR repertoire and cell state to provide effective immunity against pathogens^1,2^. However, it remains unclear how differences in the TCR contribute to heterogeneity in T cell state at the single cell level because most analysis of the TCR repertoire has, to date, aggregated information from populations of cells. Single cell RNA-sequencing (scRNA-seq) can allow simultaneous measurement of TCR sequence and global transcriptional profile from single cells. However, current protocols to directly sequence the TCR require the use of long sequencing reads, increasing the cost and decreasing the number of cells that can be feasibly analyzed. Here we present a tool that can efficiently extract TCR sequence information from standard, short-read scRNA-seq libraries of T cells: TCR Reconstruction Algorithm for Paired-End Single cell (TRAPeS). We apply it to investigate heterogeneity in the CD8^+^ T cell response in humans and mice, and show that it is accurate and more sensitive than previous approaches^3,4^. We applied TRAPeS to single cell RNA-seq of CD8^+^ T cells specific for a single epitope from Yellow Fever Virus^5^. We show that the recently-described "naive-like" memory population of YFV-specific CD8^+^ T cells have significantly longer CDR3 regions and greater divergence from germline sequence than do effector-memory phenotype CD8^+^ T cells specific for YFV. This suggests that TCR usage contributes to heterogeneity in the differentiation state of the CD8^+^ T cell response to YFV. TRAPeS is publicly available, and can be readily used to investigate the relationship between the TCR repertoire and cellular phenotype.

## MAIN

The population of antigen-specific CD8^+^ T cells formed in response to infection or vaccination is highly heterogeneous in terms of function and phenotype ^6,7^. Efforts to deconvolve this cellular heterogeneity have used flow cytometry, mass spectrometry, and more recently, single-cell RNA-sequencing ^8^. These approaches have identified a reliable set of phenotypic markers that can classify antigen-specific T cells into a large number of subsets, and distinguish them from antigen-naive T cells. However, recent work also suggests that some antigen-experienced CD8^+^ T cells can have a naive-like phenotype ^5,9,10^. The cellular heterogeneity in the T cell compartment is thought to arise from different exposure to differentiation cues such as antigen dose, duration of contact, and cytokines. How the TCR sequence expressed by each T cell contributes to that cellular heterogeneity is not fully understood.

The T cell receptor is a heterodimer of two chains - alpha and beta, each consisting of three types of genomic segments - variable (V), joining (J) and constant (C) (the beta chain includes an additional short diversity (D) segment; Methods) ^2^. The V and J segments are selected out of a pool of several dozen loci encoded in the germline genome, through a recombination process. The diversity of the TCR repertoire (estimated at ∼10^7^ in humans ^2^) is further enhanced by random insertions and deletions into the complementary determining region 3 (CDR3) – the junction between the V and J segments, which largely determines the ability of the cell to recognize specific antigens. However despite this diversity, some T cell responses can include TCRs that are identical between individuals - known as “public” clonotypes, while other T cell responses use TCRs that are unique to each individual (“private” clonotypes). Previous studies have shown that these public clonotypes tend to appear at a higher frequency and have a shorter CDR3 region, possibly as a result of a more efficient recombination process ^2,11^.

Unlike analysis of the cell state, the clonal diversity of the TCR repertoire has to date been studied mostly in aggregated samples from pools of T cells ^2,12,13^ rather than individual cells. This approach has two significant limitations: (1) since each chain of the TCR (alpha, beta) is a separate transcript, it cannot determine which chains are co-expressed in the same cell, leading to a partial view of the TCR identity; (2) the sequence of the TCR and the global transcriptional state of cell that expresses it cannot be simultaneously determined. Some studies have profiled TCR use in single cells, but these studies were limited in the number of transcripts that were quantified ^12,14^.

Single cell RNA-seq can generate full-length sequence information for many transcripts in individual cells including the alpha and beta chains of the TCR. However, standard methods to map sequence fragments to the genome ^15^ cannot be directly used for reconstructing and estimating the abundance of TCRs because of the highly variable nature of the CDR3 regions. One approach to address this challenge is to rely on scRNA-seq with long sequencing reads (>100bp), which can cover the entire CDR3 region along with the flanking V and J sub-segments ^3^. The underlying TCR (along with the junctional diversification events) can then be identified using methods similar to TCR-seq population repertoire analysis ^2,16^. However, sequencing with long reads is costly and time consuming, thus a method to successfully reconstruct TCRs from shorter, paired-end reads is desirable. Another approach, de-novo transcriptome assembly ^3,4^, has only been tested on long (>100bp paired-end) RNA-seq libraries and does not make sufficient use of the information we have about the CDR3 flanking sequences. Furthermore, de-novo assemblers were designed with a very large input set and long read length in mind. More accurate yet computationally intensive algorithms are feasible and can be applied to the smaller problem of reconstructing only the TCR (rather than the whole transcriptome).

Here, we provide an approach to address these problems. First, we present TRAPeS, a software tool capable of accurately reconstructing TCR from paired-end sequencing libraries of single cells, even at short (25bp) read length. TRAPeS makes use of the conserved genomic information of the flanking regions and applies a reconstruction scheme that leads to marked increase in sensitivity. Second, we demonstrate how simultaneous analysis of TCR properties and global expression profiling in individual cells helps relate specific TCR properties such as CDR3 length to heterogeneity of T cell state among CD8^+^ T cells that respond to YFV.

TRAPeS starts by recognizing putative pairs of V and J segments that flank the CDR3 region, using genome alignment ^17^ (Figure 1A, top; see Methods for a complete description of the algorithm). It then identifies the set of unaligned reads that may have originated from the CDR3 region, taking the unmapped mates of reads aligned to the putative V-J segments or to the constant (C) segment (Figure 1A, middle). Next, it uses an iterative dynamic programing scheme to piece together the putative CDR3 reads, gradually extending the CDR3 reconstruction on both ends (V and J) until convergence (Figure 1A, bottom). Finally, after the TCR chain has been reconstructed, TRAPeS determines whether it is productive (i.e., has an in-frame CDR3 without a stop codon) and determines its exact CDR3 sequence, based on the criteria established by the international ImMunoGeneTics information system (IMGT) ^18^. For each cell, TRAPeS outputs a set of reconstructed TCR transcripts (from both chains), along with their complete sequence, an indication of whether or not they are productive, and the number of reads mapped to them. In some cases multiple reconstructions can be generated for the same cell. This primarily happens as some V and J segments have very similar sequences, resulting in several possible V-J pairs with an identical CDR3 reconstruction. In such cases, we report all V-J pairs, while ranking the putative TCR transcripts in accordance to their estimated expression levels. The average running time of TRAPeS on a Human single cell library with an average two million reads per cell is less than two minutes per cell on a standard machine (Figure S1).

**Figure 1:**
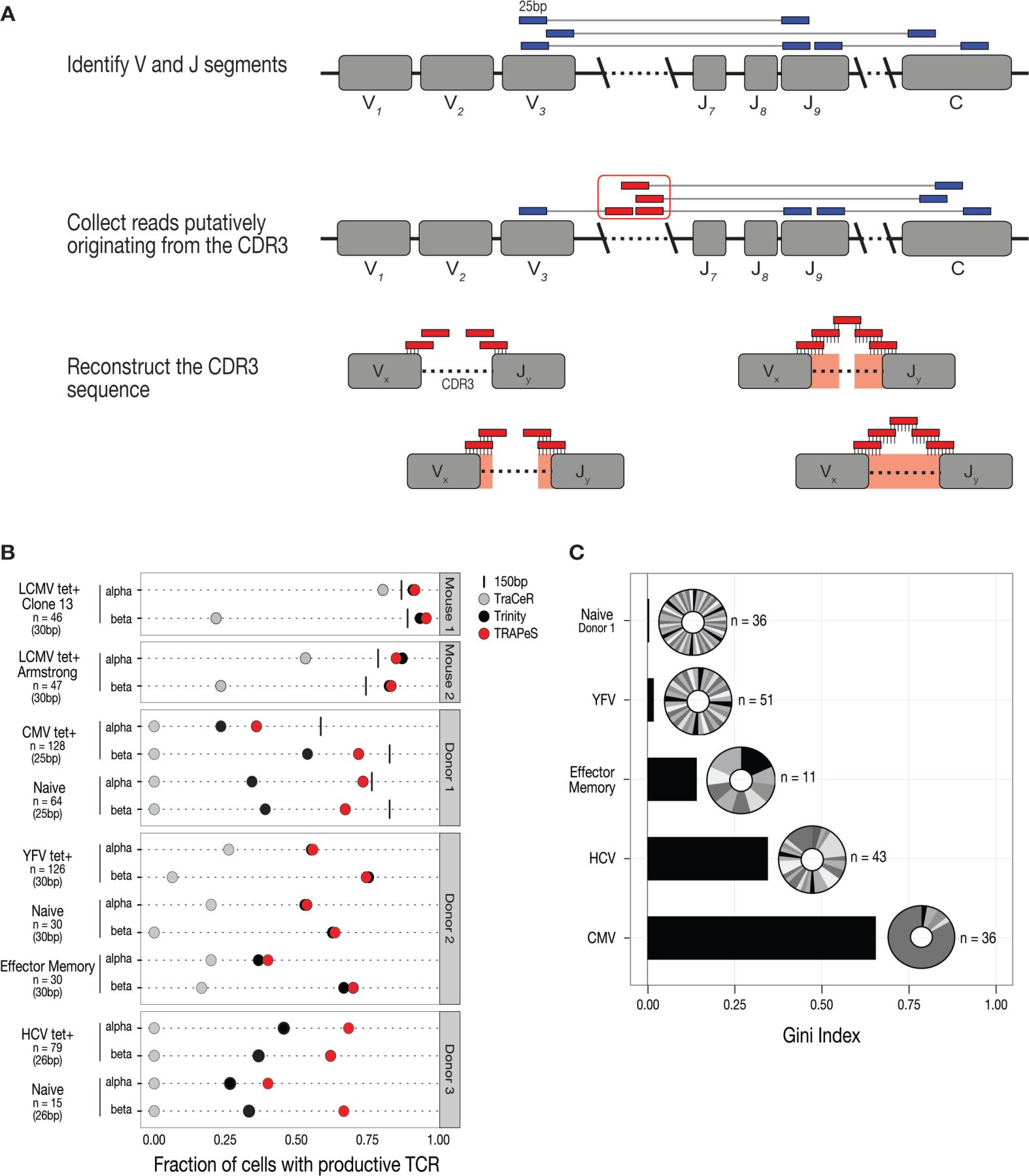
TRAPeS - An algorithm for TCR reconstruction in single cell RNA-seq. **a)** Illustration of the TRAPeS algorithm. First, the V and J segment are identified by searching for paired reads with one read mapping to the V segment and its mate mapping to the J segment. Then, a set of putative CDR3-originating reads is identified as the set of unmapped reads whose mates map to the V, J and C segments. Finally, an iterative dynamic programming algorithm is used to reconstruct the CDR3 region. **b)** Validation of the TCR reconstruction. Success rates for reconstruction of productive CDR3 in various CD8^+^ T cell data sets. Each line depicts the fraction of cells with a productive alpha or beta chain in a given data set with each one of the following methods - 150bp sequencing (black line), short paired-end data reconstructed using TRAPeS (red), TraCeR (light grey) or Trinity (black). **c)** Single cell RNA-sequencing captures a variety of clonal responses. Bars represent the Gini coefficient of each human CD8^+^ T cell data set. The Gini coefficient can range from zero (a complete heterogeneous population) to one (a complete homogeneous population). Pie charts represent the distribution of clones in each population.

We applied and tested TRAPeS to scRNA-seq data from a range of CD8^+^ T cell responses (Methods, Figure 1B). These data sets were selected to include both mouse and human CD8^+^ T cells as well as those expected to have a range of TCR complexities (Figure S2). In mice, we used the lymphocytic choriomeningitis virus (LCMV) infection model, and profiled CD8^+^ T cells responding to either acute or chronic infection (using the Armstrong and Clone 13 strains of LCMV, respectively). In healthy human subjects we profiled naive CD8^+^ T cells, effector memory CD8^+^ T cells, and antigen-specific CD8^+^ T cells elicited by CMV infection; vaccination with the live attenuated yellow fever virus infection (YFV-17D) ^19^; or by vaccination with adenoviral and modified vaccinia Ankara vectors encoding HCV proteins ^10,20^. We sorted up to 128 single CD8^+^ T cells from each dataset, and generated scRNA-seq libraries with short (25-30bp) paired-end reads as previously described ^21,22^ and observed good quality metrics using previously used measures ^1^ (Methods).

We began by evaluating the accuracy of TRAPeS by comparing its output with that from directly sequencing the TCR sequence using long reads (in which reconstruction is not required). To that end, we sequenced libraries of epitope-specific cells for Clone 13, Armstrong and CMV, and naive T cells from the CMV donor with both short (25-30bp) paired-end and 150bp single-end sequence reads (Figure 1B). TCR sequences identified by TRAPeS were almost perfectly consistent with those produced based on the long read data (Figure S3A), indicating a high level of specificity.

Next, we compared TRAPeS to TraCeR ^3^ - a recently described algorithm for TCR reconstruction that is built upon Trinity ^4^, a de-novo transcriptome assembly tool. We found that the sensitivity of TRAPeS was markedly higher (Figure 1B, Figure S3B). TRAPeS successfully reconstructed an average of 60% productive alpha chains and 73% productive beta chains. In contrast, TraCeR resulted in no reconstruction for the 25bp paired-end datasets, and was able to determine CDR3 region in only 40% of cells with productive alpha and 14% of cells with productive beta for the 30bp sequencing. This is likely due to TraCeR’s requirement for seed *k*-mer length (25nt) that is unsuitable for short reads. Thus, we also ran Trinity on our set of putative CDR3-originating reads, using a *k* value of 13. This resulted in an increased sensitivity, albeit still substantially less than TRAPeS (Figure 1B and Figure S3B). Importantly, the rate of successful reconstruction of productive chains with TRAPeS using short reads (25-30bp paired-end), is comparable to that of TraCeR ^3^ using long reads (100bp, paired-end). This is evident by application of TRAPeS to a trimmed version of the data used by Stubbington et al. ^3^(Figure S4), or by a comparison of reconstruction rates after applying a similar cell-quality filtering scheme (Figure S5).

Next, we investigated the clonality of the TCR repertoire measured by TRAPeS among the human CD8^+^ T cells (Figure 1C), using the Gini Index, a clonality measure ^23^ ranging from zero (i.e. no two cells share the same TCR) to one (i.e. all cells are from the same clone; Methods). As expected, the naïve population had a Gini index of zero, indicating that each naive CD8^+^ T cell expressed a unique TCR. The CMV-specific CD8^+^ T cell population had a high Gini index (with 83% of CMV-specific CD8^+^ T cells with reconstructed alpha and beta chains originated from a single clone), indicating a high degree of oligoclonality as previously described ^24,25^. In contrast, CD8^+^ T cells elicited by YFV or HCV vaccines showed much greater heterogeneity in TCR repertoire, consistent with a more limited, rather than persistent, exposure to antigen ^10,26–29^.

In order to determine the relationship between TCR use and CD8^+^ T cell state, we focused on CD8^+^ T cells from healthy donors (YFV and CMV peptide-specific, naive or effector memory) to avoid introducing additional complexity from chronic infection. To identify groups of cells with similar expression profiles, we used SC3 ^30^, a robust clustering method for sparse datasets, to identify subpopulations of cells (Figure S6, Methods) which we then visualized using t-SNE ^31^ (Figure 2A). We found three clusters of cells: one that contained all CMV-specific cells (Figure 2A, purple symbols); one that contained all effector memory cells (blue symbols); and one that contained all naive CD8^+^ T cells (green symbols). In contrast to these discrete groupings, we observed that YFV-specific CD8^+^ T cells were split between two clusters: one containing effector memory CD8^+^ T cells and one containing naive CD8^+^ T cells.

**Figure 2:**
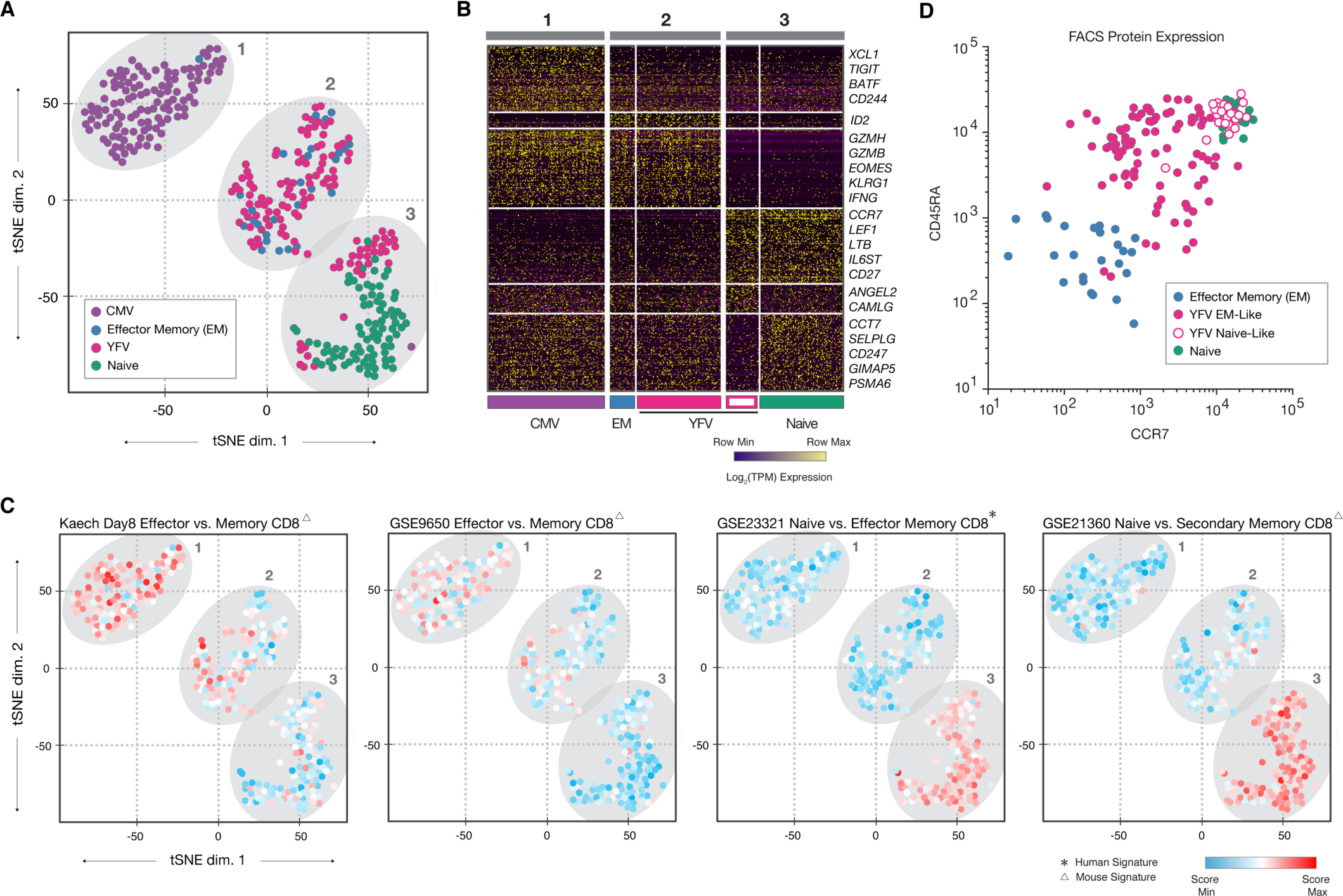
Transcriptome analysis reveals distinct subpopulation of YFV-specific cells exhibiting a naive-like profile. **a)** t-SNE projection of 353 CMV-specific, Effector Memory, YFV-specific, and Naïve cells, using normalized TPM values of 10827 transcripts. Ellipses indicate three distinct spatial clusters. A discrete subset of YFV-specific cells cluster with Naive. **b)** Genes differentially expressed between relevant phenotypic groups. YFV-specific cells were classified as effector memory-like or naive-like using SC3, a non-spatial consensus clustering approach (Figure S6). **c)** t-SNE projections, each cell colored by relative signature score. Shown are two signatures from the ImmuneSigDB distinguishing CMV-specific from YFV-specific cells, and two signatures distinguishing Naïve or YFV-specific naive-like cells from Effector memory, CMV-specific and YFV-specific effector memory-like populations. **d)** FACS protein expression of CCR7 and CD45RA surface molecules from index sort of Effector Memory, YFV-specific effector memory-like, YFV-specific naive-like, and Naïve cells.

Differential gene expression analysis between cell clusters revealed transcripts consistent with the known patterns of gene expression in antigen-experienced or naive CD8^+^ T cells (Figure 2B). CMV-specific CD8^+^ T cells expressed effector molecules and transcription factors characteristic of antigen experienced cells (e.g., Granzyme B, *PRDM1*), which were not detected in naive cells. Naive CD8^+^ T cells expressed canonical markers of the naive state (*CCR7*, *SATB1*, *LEF1*) that were absent in CMV-specific and effector memory CD8^+^ T cells. The expression of these genes in YFV-specific CD8^+^ T cells was consistent with the cluster in which they were associated, with those in the naive cluster expressing minimal Granzyme B or *PRDM1*, but showing robust expression of *CCR7*, *SATB1*, and *LEF1* (Figure S7).

To identify broader patterns of transcriptional signatures, we applied FastProject ^32^ - a software tool that enables the expression of gene sets of interest to be quantified in transcriptional profiles of single cells (Methods). We surveyed the representation of a collection of gene sets, from the C7 (ImmuneSigDB) ^33^ collection of MSigDB ^34^ corresponding to cell states and perturbations of CD8^+^ T cells. We found significant upregulation of multiple gene sets corresponding to naive CD8^+^ T cells (K-S test FDR-adjusted p-value<0.01) in naive-like cluster (cluster 3) compared to the other two clusters. Consistent with this, we found significantly greater upregulation of effector signatures in clusters 1 and 2 compared with the other clusters (FDR-adjusted p-value<0.01; Figure 2C).

To confirm these patterns of transcript abundance at the protein level, we compared flow cytometry data for a set of surface markers acquired at the time of sorting (Methods) with transcript abundance in the each cell (Figure 2D). Consistent with the gene expression profiles, we observed that YFV-specific CD8^+^ T cells in the naive-like cluster (open symbols) showed higher protein levels of CCR7 and CD45RA than those in the effector memory cluster (purple symbols). Thus, single-cell analysis shows that CD8^+^ T cells specific for the same peptide epitope from YFV are heterogeneous and includes both effector-memory and naive-like gene expression profiles, as has been reported previously for cells analyzed at the bulk level ^5,9,35^.

We reasoned that differences in TCR might contribute to the heterogeneous differentiation of CD8^+^ T cells following YFV vaccination. To that end, we evaluated a number of properties to characterize each reconstructed TCR, and asked whether any differed between naive-like and effector memory-like YFV-specific CD8^+^ T cells. Naive-like and effector memory-like YFV-specific CD8^+^ T cells were indistinguishable (p-value>0.05) in terms of TCR transcript expression, hydrophobicity of the CDR3 region and tetramer binding (Methods). However, we found that the CDR3 sequence was significantly longer in YFV-specific CD8^+^ T cells with a naive-like state compared with those with an effector memory profile for both alpha and beta chains (Figure 3A, K-S test p-value 0.038 and 0.027 for alpha and beta chains, respectively).

**Figure 3:**
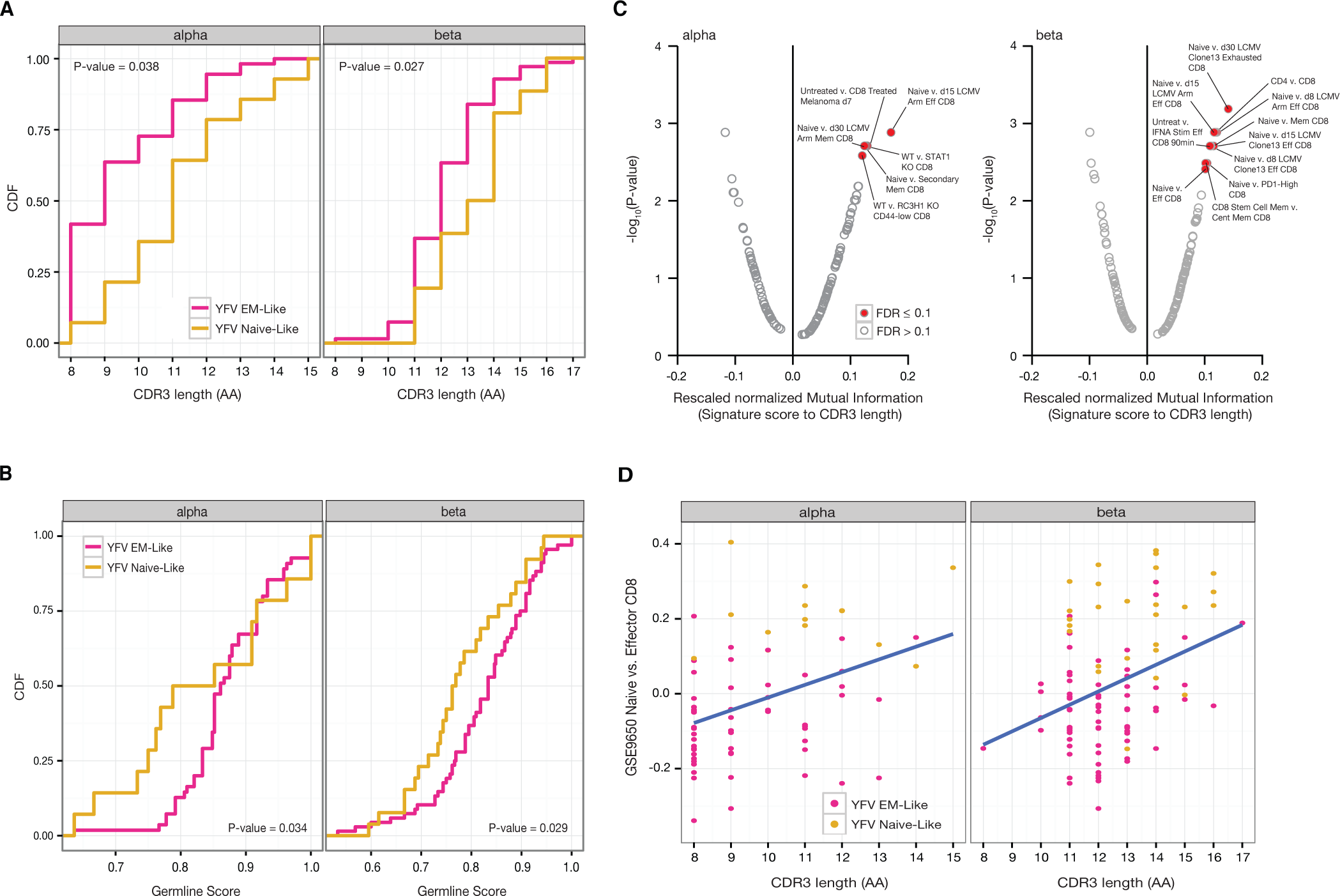
YFV-specific subpopulations display different TCR structure. **a)** YFV-specific naive-like cells tend to have longer CDR3. Distribution of the YFV-specific effector memory-like and naive-like CDR3 lengths in both alpha (left) and beta (right) chains. P-values were calculated with K-S test. **b)** Differences between naive-like and effector memory-like CDR3 lengths are due to added nucleotides. Distribution of the YFV-specific effector memory-like and naive-like CDR3 germline scores, defined as the number of nucleotides in the CDR3 encoded by the V, D or J segments divided by the total number of nucleotides in the CDR3, for both alpha (left) and beta (right) chains. P-values were calculated with K-S test. **c**) Signature analysis reveals significant correlation between CDR3 length and cell state. The plot depicts the rescaled normalized mutual information score between CDR3 length and transcriptional signatures of CD8^+^ T cells from ImmuneSigDB. Signatures identified as statistically significant using a permutation test (FDR-adjusted p-value<0.1) are highlighted in red. **d)** YFV-specific cells with long CDR3 tend to have a higher transcriptomic naive signature than cells with short CDR3. Plot represents the score of each cell for a transcriptional signature of a naive vs. effector CD8^+^ T cell state. A high signature score means that a cell has higher expression of naive signature genes compared to effector signature genes.

We next evaluated the germline score of CDR3 regions in YFV-specific CD8^+^ T cells, a measure of the contribution of germline nucleotides to the CDR3 region. The germline score is defined as the ratio between the number of nucleotides in the CDR3 that originate from the germline (V, D, J segments) to the total number of nucleotides in the CDR3 ^16^ (Methods). Consistent with the differences in the CDR3 length, we found that naive-like YFV-specific CD8^+^ T cells had a significantly lower germline score in both alpha and beta chains than did effector memory-like cells (Figure 3B, K-S test p-value of 0.034 and 0.029 for alpha and beta chains, respectively), suggesting that generating the CDR3 region of these TCRs involved a greater degree of nucleotide addition/subtraction.

To further characterize the relationship between CDR3 length and cellular state in YFV-specific CD8^+^ T cells, we identified CD8^+^ transcriptional signatures (extracted from ImmuneSigDB ^33^ and scored with FastProject ^32^, as above) that correlated with CDR3 length across all YFV-specific CD8^+^ T cells (Methods). Of all signatures evaluated, we found that only naive CD8^+^ T cell signatures showed a significant positive correlation with CDR3 length (FDR-adjusted p-value<0.1; Figures 3C-3D). Previous work has suggested that YFV-specific CD8^+^ T cells with a naive-like phenotype include those with a stem-cell memory (Tstem-memory) differentiation state. We found that Tstem-memory signatures were more enriched in naive-like YFV-specific CD8^+^ T cells than in effector memory YFV-specific CD8^+^ T cells (Figure S8). However, the enrichment for these signatures was equivalent between naive-like YFV-specific and phenotypically naive CD8^+^ T cells, making it difficult to discern whether these cells manifest a specific stem-cell-like state. Our results, however, show that heterogeneity in the differentiation state of CD8^+^ T cells responding to a single epitope of YFV is strongly associated with the CDR3 length.

TRAPeS enables the analysis of TCR clonality in scRNA-seq profiles using short sequence reads. Other methods of direct TCR sequencing ^2^ or reconstruction ^3^ required long sequence reads, which substantially increase the per-cell cost of single cell profiling. As single-cell RNA-seq technologies move towards massively parallel scale, long-read sequencing is likely to become unfeasibly expensive, making approaches such as TRAPeS critical for studies of TCR use in single cells.

We applied TRAPeS to short-read sequencing data from single human CD8^+^ T cells to discover a new association between the differentiation state of antigen-specific CD8^+^ T cells and the CDR3 length of the TCRs that they express. Long CDR3 lengths have been associated with private clonotypes, which in turn may reflect low precursor frequency within the naive T cell pool ^2,11^. We therefore speculate that within a population of naive T cells capable of recognizing a specific antigen, those that exist at low frequency may enter the T cell response later than more abundant precursors, resulting in an altered differentiation state compared to those that existed at a higher precursor frequency. Alternatively, a greater degree of cross-reactivity in T cells with short CDR3 regions may result in more repeated TCR stimulation, leading to the difference in T cell phenotype we observe. More generally, we anticipate that TRAPeS will facilitate broad efforts to determine the relationship between T cell state and TCR sequence in the immune response to vaccination and cancer.

## METHODS

### TRAPeS

The TRAPeS algorithm has 4 main steps, each applied separately to the alpha and beta chains:

1. **Identifying putative pairs of V and J segments**. In order to recognize the V and J segments of the TCR, TRAPeS takes as input the alignment of the RNA-seq reads to the genome. TRAPeS searches for a paired-end read where one mate maps to a V segment while the other mate is mapped to a J segment, and takes those V-J pairs as putative candidates for the CDR3 reconstruction. In a case where there is no such pairing, TRAPeS takes all possible V-J combinations of V and J segments that have V-C and J-C pairing (i.e. reads where one mate maps to V or J and the other mate maps to the C segment). We note that reads are not successfully aligned to D segments of the beta chain due to their short length, thus for the beta chain reconstruction of the CDR3 includes reconstruction of the D segment sequence.
2. **Collecting putative CDR3-originating reads**. TRAPeS finds the putative CDR3-originating reads by taking all the unmapped reads whose mates map to the V/J/C segments. In addition, since the first step of the CDR3 reconstruction includes alignment to the germline V/J sequences (see below), TRAPeS also collects the reads that map to the V and J segments.
3. **Reconstructing the CDR3**. Using an iterative dynamic programing algorithm, we extend the V and J regions. At the initial step, we take the ends for the V and J segments closest to the CDR3 (3’ of the V segment and 5’ of the J segment). The number of initial bases is a parameter that can be tuned, set by default to min(length(V), length(J)). In each iteration, we align all the reads to the V and J segments separately with the Needleman-Wunsch algorithm, using the following scoring scheme: +1 for a match, -1 for a mismatch, -20 for gap opening and -4 for gap extension. In addition, we don’t penalize for having the read “flank” the V and J toward their 3’ and 5’, respectively. Next, we take all the reads that aligned to the V and J segments above a certain score threshold, and build the “extended” V and J sequences based on the reads. For each position, we take the base that appears in most reads as the chosen base for this position. This way, we extend the V and J regions in each iteration and also correct for mutations or SNPs in the known genomic V and J segments. For this work, we used a threshold score of 21 for the alignment of the reads. However, in some cases a lower threshold was required, thus if no sequence was reconstructed we run TRAPeS with a scoring threshold of 15.
4. **Separating similar TCRs and determining chain productivity**. Since some V and J segments have similar sequence, reads can be mapped to several segments, creating few similar putative V-J pairs. TRAPeS takes all those possible pairing and attempts to reconstruct the CDR3 region for all pairs. After reconstruction, TRAPeS runs RSEM ^15^ on all reconstructed TCRs and the set of reads used as input (and their mates) in order to determine the relative abundance of each TCR and rank the TCRs based on the relative abundance. Next, TRAPeS determines if the TCR is productive: V and J segments are in the same reading frame and the CDR3 does not contain a stop codon. TRAPeS outputs all possible reconstructions. For this paper we used the productive chain with the highest expression as the TCR sequence for each cell.

TRAPeS is implemented in python. To increase performance, the CDR3 reconstruction using the dynamic programming algorithm is implemented in C++, and uses the seqan package ^36^. TRAPeS is freely available and can be downloaded in the following link: https://github.com/YosefLab/TRAPeS

### Single cell sorting

*Mouse LCMV Experiments:*Female C57BL/6 mice (The Jackson Laboratory), aged 7 weeks, were infected with 2x10^5^ plaque forming units (PFU) LCMV Armstrong intraperitoneally i.p. or 4x10^6^ PFU LCMV Clone 13 i.v. LCMV viruses were a generous gift from Dr. E John Wherry (University of Pennsylvania, Perelman School of Medicine). Peripheral blood was obtained from the mice at day 7 post infection (p.i.) and lymphocytes were enriched using LSM density centrifugation. Cells were prestained with a near-IR fixable live/dead marker (Life Technologies, cat# L34976) and an APC-conjugated dextramer reagent for gp33 (Immudex, cat# A2160-APC) according to manufacturer recommendations. The cells were then stained with the following antibodies: FITC 2B4 (BioLegend, cat# 133504), PerCP-Cy5.5 CD44 (BioLegend, cat# 103032), PE KLRG1 (BioLegend, cat# 138408), PE-Cy7 PD1 (BioLegend, cat# 135215), BV421 CD127 (BioLegend, cat# 135024), BV510 CD8A (BioLegend, cat# 100752).

*Human CMV Experiments (Donor 1):*Blood samples were obtained from a donor with a detectable NLV-specific CD8^+^ T cell response. Lymphocytes were enriched via Ficoll gradient and prestained with a fixable live/dead marker (Life Technologies, cat# L34976) and an APC-conjugated dextramer reagent (Immudex, cat# WB2132-APC). The cells were then stained with the following antibodies: FITC CD8A (BioLegend, cat# 300906), PerCP-Cy5.5 CCR7 (BioLegend, cat# 353220), PE CD3 (BioLegend, cat# 317308), BV605 CD45RA (BioLegend, cat# 304133).

*Human YFV Experiment (Donor 2)*: A healthy volunteer was vaccinated with a single dose (0.5 ml containing at least 10^5^ PFU) of 17D live-attenuated yellow fever vaccine strain administered subcutaneously. Seroconversion after vaccination was confirmed by assaying the neutralizing antibody titers for YF-17D (data not shown). A whole blood sample was obtained 9 months post-vaccination and lymphocytes were enriched from whole blood via Ficoll gradient centrifugation and a CD8 negative selection magnetic bead kit. Cells were prestained with a live/dead marker (Life Technologies, cat# L34976) and an APC-labeled tetramer reagent (NS4B 214–222 LLWNGPMAV, kindly provided by Dr. Rafi Ahmed). The cells were then stained with the following antibodies: FITC CD8A (BioLegend, cat# 300906), PE CXCR3 (BioLegend, cat# 353705), PE-Cy7 CCR7 (BioLegend, cat# 353226), BV421 IL2Rb (BioLegend, cat# 339009), BV510 CD3 (BioLegend, cat# 317332), BV605 CD95 (BioLegend, cat# 305627), BV780 CD45RA (BioLegend, cat# 304140).

*Human Hepatitis C Experiment (Donor 3):* Patient 355 (59yr old Male, infected with genotype 1a HCV, baseline viral load 467,000 IU/ml) received a prime vaccination of ChAd3-NSmut (2.5x10^10^ viral particles) and an MVA-NSmut (2x10^8^ plaque forming units) boost vaccination 8 weeks later. PBMC were collected 14 weeks post-boost vaccination for assessment of single cell gene expression ^20^. PBMC were thawed and prestained with a live/dead marker (Life Technologies, cat# L34976) and a PE-conjugated pentamer reagent (PE-labeled HCV NS31406–1415 (KLSALGINAV; HLA-A*0201)). The cells were then stained with the following antibodies: FITC 2B4 (BioLegend, cat# 329505), PerCP-eFluor 710 LAG3 (eBioscience, cat# 46-2239), PE-Cy7 CCR7 (BioLegend, cat# 353226), APC CD39 (BioLegend, cat# 328209), BV421 PD1 (BioLegend, cat# 329919), BV510 CD3 (BioLegend, cat# 317332), BV605 CD8A (BioLegend, cat# 301040), BV780 CD45RA (BioLegend, cat# 304140).

The relevant institutional review boards approved all human subject protocols, and all subjects provided written informed consent before enrollment.

*Single cell sorts:*All single cell sorts were performed on a BD Aria II with a 70um nozzle. Cells were sorted into 5μL of Qiagen RLT plus 1% beta-mercaptoethanol v/v. Immediately following sorting, plates were sealed, vortexed on high for 30 seconds, and spun at 400g for 1 minute prior to flash freezing on dry ice. Samples were stored at -80C until library preparation.

### RNA sequencing

Single cell lysates were converted to cDNA following capture with Agencourt RNA Clean beads using the SmartSeq2 protocol as previously described ^21^. The cDNA was amplified using 22-24 PCR enrichment cycles prior to quantification and dual-index barcoding with the Illumina Nextera XT kit. The libraries were enriched with 12 cycles of PCR, then combined in equal volumes prior to final bead cleanup and sequencing. All libraries were sequenced on an Illumina HiSeq 2500 or NextSeq by either single-end 150bp reads or short paired-end reads using the following read lengths: Mouse samples - 30bp, Human donor 1 - 26bp for read 1 and 25bp for read 2, Human donor 2 - 30bp, human donor 3 - 26bp.

### Preprocessing and Normalization of scRNA-seq data

Low quality bases were trimmed with trimmomatic ^37^ using the following parameters: LEADING:15, TRAILING:15, SLIDINGWINDOW:4:15, MINLEN:16. Trimmed reads were then aligned to the genome (hg38 or mm10 for human or mouse samples, respectively) with TopHat2 ^38^ for TCR reconstruction, and aligned to the transcriptome with RSEM ^15^ for transcriptome quantification.

For transcriptome analysis of the human CMV and YFV donors (donors 1 and 2), low quality cells were filtered out prior to normalization. Cells were filtered out if their read depth was less than 1 million pairs or if the cell expressed less than 20% of all expressed transcripts, where a transcript was considered expressed if it had a TPM value of >10 in at least 10% of cells.

Normalization of TPM values was done with our newly developed normalization framework (SCONE ^39^). First, each sample is scaled with the DEseq ^40^ scaling factor to account for differences in sequencing depth. Then, to remove unwanted variance from the data we ran RUVg ^41^ with k=1. In order to run RUVg, a list of genes that are constant across conditions should be provided. To find constant genes across the specific conditions that were tested in this paper, we also sequenced populations of 100 cells of naive CD8^+^ T cells from donor 1 and CMV-specific effector memory CD8^+^ T cells, as well as populations of 50 cells of naive CD8^+^ T cells from donor 2 and YFV-specific effector memory CD8^+^ T cells. We ran DESeq2 ^42^ on those samples and defined the set of constant genes as the genes that showed no change (FDR-adjusted p-value > 0.98 and absolute log fold change < 0.2) across all pairwise comparisons (naive vs. all effector memory cells, naive vs. CMV-specific effector memory, naive vs. YFV-specific effector memory and CMV-specific effector memory vs. YFV-specific effector memory), resulting in a total of 373 genes.

### Gini coefficient calculation

For each population, cells were considered from the same clone if the had identical CDR3 sequences of both alpha and beta chains. Cells with only one reconstructed chain were excluded from this analysis. The number of cells for each clone was counted and the Gini coefficient was calculated by using the Gini command in R from the “ineq” package ^43^.

### Inference of cell clusters, visualization and differential expression analysis

For cluster inference in the YFV + CMV human data, we defined an expression matrix consisting of normalized TPM values of 353 cells by 10827 transcripts (expressed at a level of >= 5 TPM in at least 1% of cells). We applied the SC3 software ^30^ for clustering the cells in this matrix using default parameters.

To visualize the data, we first used the jackStraw package ^44^ to reduce the dimensionality of the data and retain only principal components (PC) that are statistically significant (p-value<10^-4^) in terms of the respective percent of explained variance. This analysis retained the first three PCs. We then applied t-SNE ^31^ with default parameters and 2000 iterations to these significant PCs, further reducing the data for visualization in two dimensions.

We used the DESeq2 package ^42^ to identify genes that are differentially expressed (DE) between the different clusters. In this application, each cluster was compared to the other two clusters, looking for genes that are differentially expressed. Genes were called as differentially expressed using an FDR-adjusted p-value cutoff of 0.05. The heatmap in Figure 2B was populated with log_2_(TPM) values for genes identified as uniquely up- or down-regulated in each of the three major phenotypic groups. Enrichment of DE genes with respect to immunological pathways was determined using a Fisher exact test (FDR-adjusted p-value<10^-3^) quantifying the significance of overlap between differential genes and signatures from the ImmuneSigDB database.

### Gene signature analysis

Cell scores were computed for each transcriptomic signature with FastProject ^32^. In short, each signature is comprised of genes that are upregulated and downregulated between two cell states. For each cell, the signature score is computed by aggregating over the weighted standardized (Z-normalized across all cells) log expression levels of the signature upregulated genes, minus the weighted standardized expression of the downregulated genes. To include only relevant signatures, we analyzed only signatures with a significant consistency score (FDR-adjusted p-value<0.05) in at least one projection. In addition, only signatures that include ‘CD8’ in their name were used for further analysis, leaving a total of 95 signatures for the YFV + CMV data sets and 154 signatures for the YFV-specific analysis.

### Characterization of TCR properties of YFV-specific cells

#### TCR expression

To compute the expression of each reconstructed TCR, we added the reconstructed sequences to the transcriptome and ran RSEM on the complete extended transcriptome, using the original sequencing data (the complete fastq files) as input. This was performed for each cell separately, i.e. for each cell only its TCR sequences were added to the transcriptome. In cases where a cell had more than one reconstructed alpha or beta chain (by having two productive chains or having one productive and one unproductive chain) they were both added to the transcriptome.

#### Germline score

Classification of each base in the CDR3 as germline (originating from the V, D, J regions) or added nucleotide was done by running the full reconstructed TCR sequences thorough IMGT/V-Quest ^45,46^. The germline score was calculated by dividing the number of nucleotides encoded by V, D, J segments by the length of the CDR3 ^16^.

#### Comparing transcriptomic signatures with TCR length

Identification of gene signatures associated with TCR length was done with the PARIS algorithm ^47^, a module in GenePattern ^48^. PARIS describes the association between each signature score and TCR length by estimating their differential mutual information. For each signature, the mutual information is computed between the TCR length and the signature, and then normalized using the joint entropy. This score is rescaled with the mean of the score of the TCR length against itself and the score of the signature against itself, resulting in a rescaled normalized mutual information (RNMI) matching score. The significance of the score is evaluated by a permutation test (performed on the TCR length) and then FDR correction.

#### Hydrophobicity

The mean hydrophobicity of each CDR3 was computed using the Kyte-Doolittle ^49^ numeric hydrophobicity scale. In order to account for CDR3 length, we also computed mean hydrophobicity for each CDR3 using a sliding window (of both size 3 and 5), taking the mean across all windows. However, the sliding window also didn’t result in significant differences between YFV-specific naive-like and YFV-specific effector memory-like cells (K-S test p-value > 0.1, data not shown).

## ACKNOWLEDGMENTS

The authors would like to thank Rama Akondy and Rafi Ahmed for providing the YFV vaccine samples and related reagents; members of the Haining and Yosef lab for input; and subjects for their participation in the studies. L.S and E.B. are funded by the Medical Research Council UK (L.S as an MRC CASE studentship). This work was supported by US National Institutes of Health grants AI090023, AI057266 and AI082630.

## AUTHORS’ CONTRIBUTION

S.A. wrote TRAPeS and performed the TCR computational analysis. K.B.Y. performed the experiments. K.B. performed the transcriptome analysis. S.D. analyzed the 150bp single-end RNA-sequencing. J.G. infected the mice with LCMV and provided peripheral blood and animals for sacrifice. U.G. stained and sorted CMV cells. L.S assisted with staining and sorting of the HCV sample. P.K. and E.B. provided the human HCV sample and had input into the manuscript. D.C.D and A.H.S helped interpret the data. W.N.H. and N.Y. conceived, designed and coordinated the study. S.A, K.B.Y, K.B, W.N.H and N.Y wrote the manuscript.

---

**Supplementary Figure 1:**
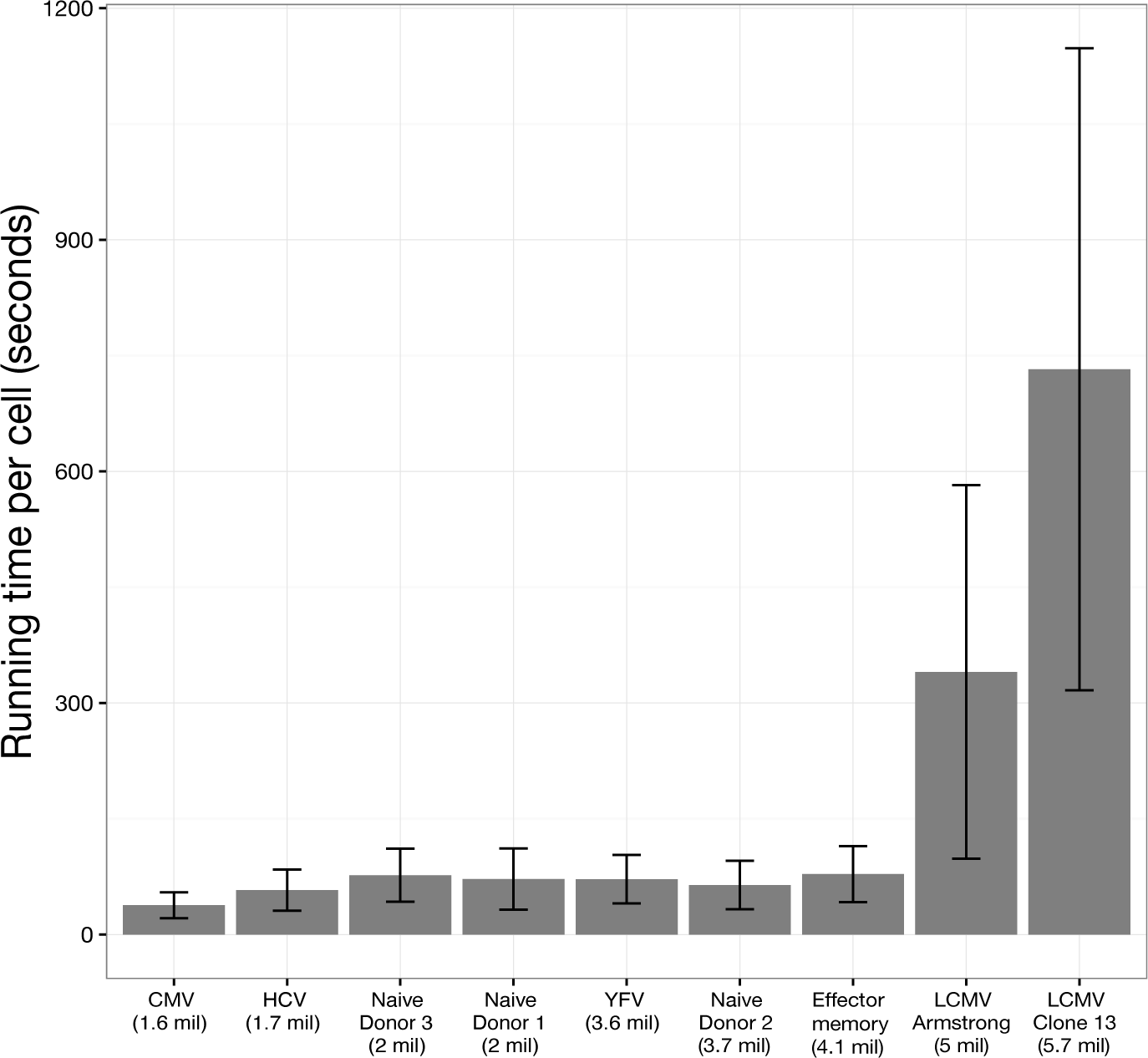
TRAPeS run times -. Average running time (in seconds) of TRAPeS per cell on all CD8^+^ T cell datasets analyzed in this study, using a single GHz processor and 8 threads. Error bars represent the standard deviation. For each dataset the average number of reads per cell is mentioned in parentheses. The increased running time in mouse samples is due to the larger sequencing depth but also due to the number of similar V and J segments which resulted in a larger number of possible V-J pairs, increasing running time.

**Supplementary Figure 2:**
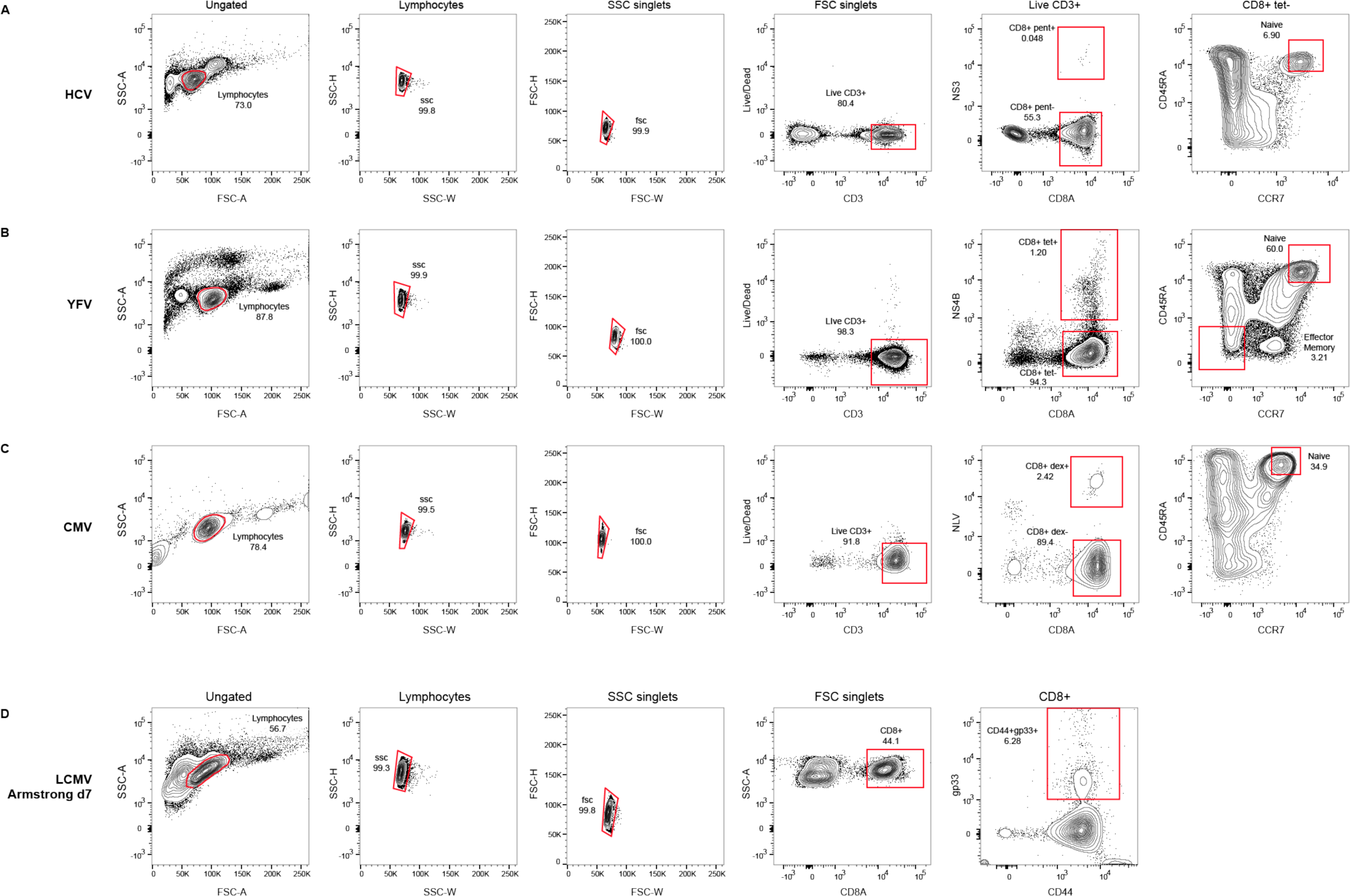
Gating strategy for CD8^+^T lymphocytes for scRNA-seq. a) Human lymphocytes were gated based on forward-scatter (FSC) and side-scatter (SSC) characteristics, then singlets were selected from SSC and FSC projections, and Live/Dead-negative CD3^+^ cells, then CD8^+^ HCV NS3 (1406-1415; KLSALGINAV; HLA-A*0201)^+^ cells were selected for HCV-specificity. CD8^+^ HCV NS3-cells were gated and then CCR7^+^ and CD45RA^+^ cells were selected to represent bulk naive CD8^+^. b) Lymphocytes were gated based on FSC and SSC characteristics, then singlets were selected from SSC and FSC projections, and Live/Dead-negative CD3^+^ cells, then CD8^+^ YFV NS4 (LLWNGPMAV)^+^ cells were selected for YFV-specificity. CD8^+^ YFV NS4-cells were gated and then CCR7^+^CD45RA^+^ and CCR7-CD45RA-cells were selected to represent bulk naive and effector memory CD8^+^, respectively. c) Lymphocytes were gated based on FSC and SSC characteristics, then singlets were selected from SSC and FSC projections, and Live/Dead-negative CD3^+^ cells, then CD8^+^ CMV NLV^+^ cells were selected for CMV-specificity. CD8^+^ CMV NLV-cells were gated and then CCR7^+^CD45RA^+^ cells were selected to represent bulk naive CD8^+^. d) Mouse lymphocytes were gated based on FSC and SSC characteristics, then singlets were selected from SSC and FSC projections, from which CD8^+^ cells, then CD44^+^ gp33^+^ cells were selected for LCMV-specificity.

**Supplementary Figure 3:**
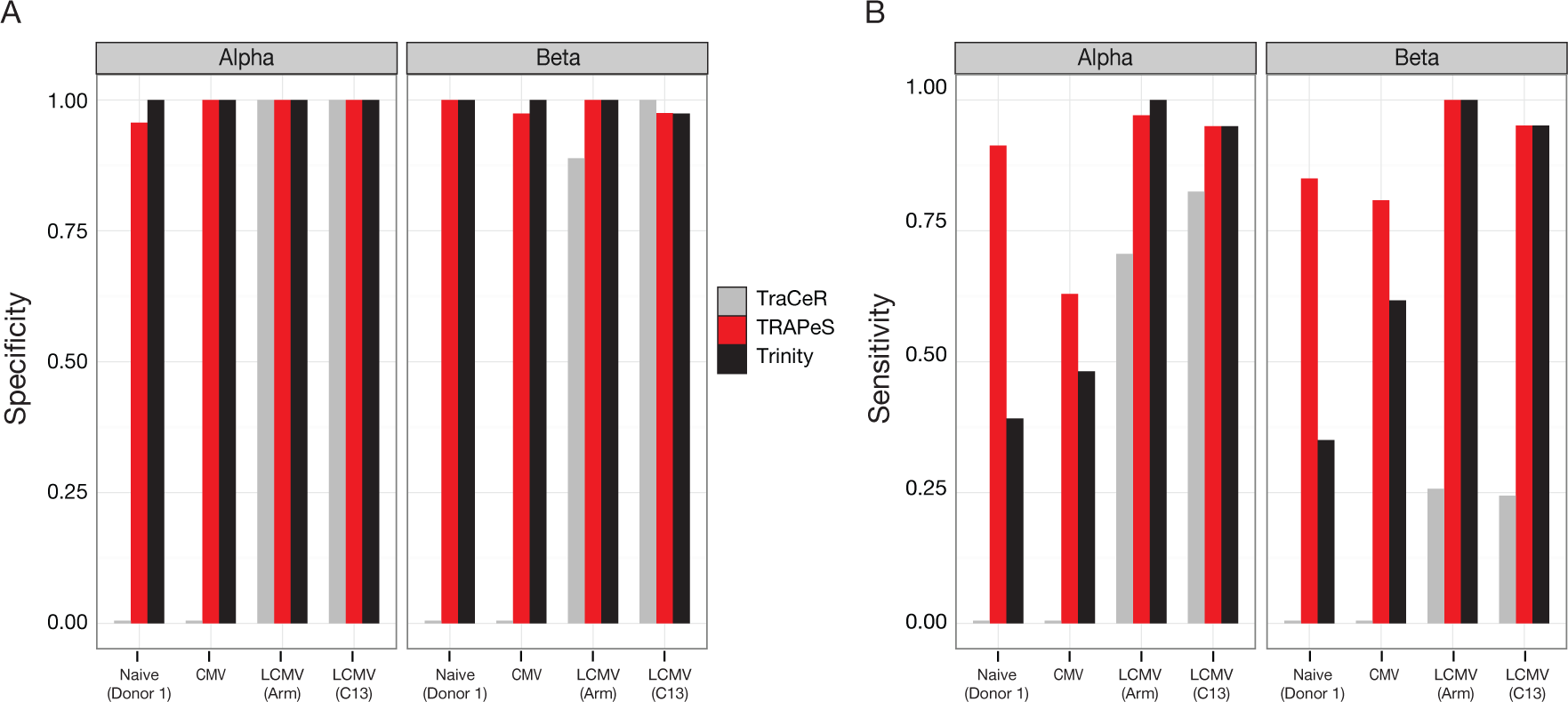
Specificity and sensitivity of TRAPeS. a) Fraction of cells with identical CDR3 sequence between 150bp data and the 25-30bp data reconstructed either by TRAPeS, TraCeR or Trinity. This was calculated as the fraction out of cells with a productive chain in both 150 and 25-30bp data. b). Same as a, except the fraction of cells is calculated out of the total number of cells that had a successful reconstruction using 150bp sequencing only.

**Supplementary Figure 4:**
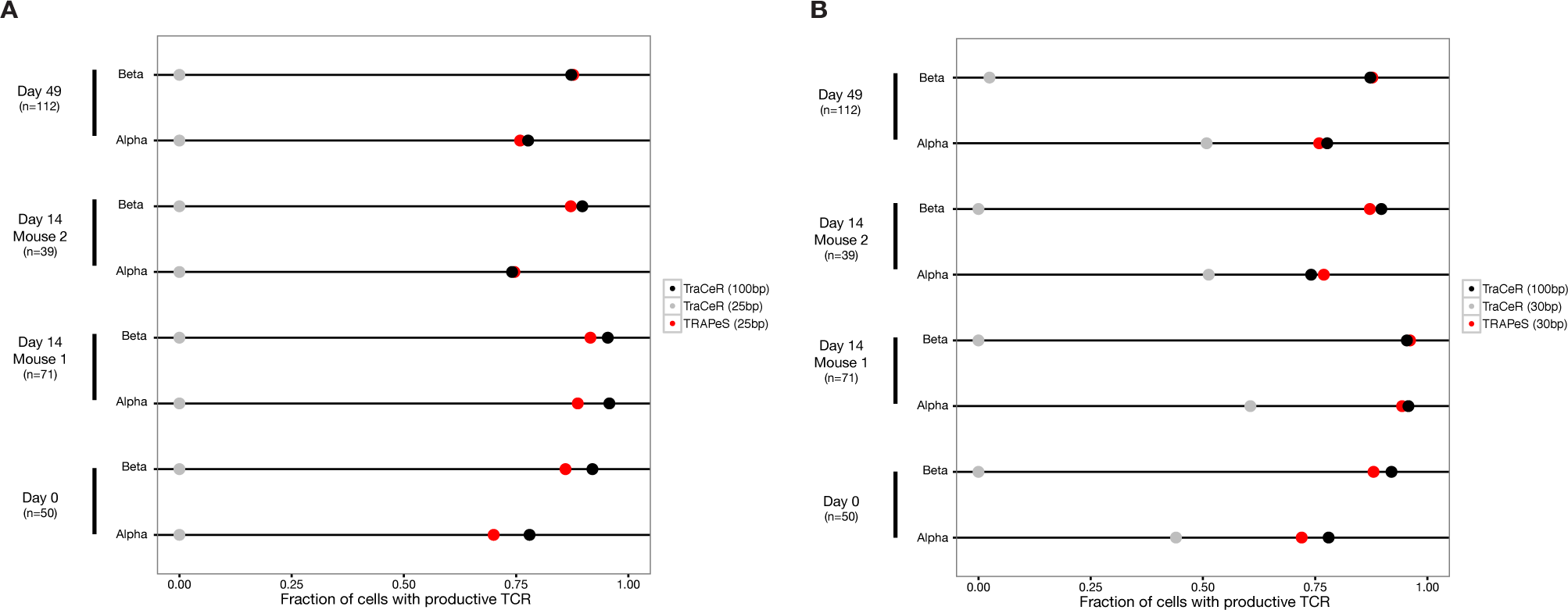
Success rates for reconstruction of productive CDR3 in the benchmark data sets used by TraCeR (Stubbington et al.) Raw single cell RNA-seq data have been downloaded as Fastq files from the Stubbington et al. study (ArrayExpress accession number E-MTAB-3857), including 272 CD4^+^ T cells from an uninfected mouse, two mice with Salmonella typhimurium infection at day 14 and one mouse at day 49 post-infection. While the original data consisted of 100bp paired-end reads, we converted it to that equivalent of “short-read” sequences by trimming each fragment to leave only the outer 25 or 30bp of each read, thus discarding 70-75% of the information. a) Each line depicts the fraction of cells with a productive alpha or beta chain in a given data set with each one of the following methods: original data (100bp paired end) with TraCeR (black); Trimmed data (taking only the outer 25bp from each read) with TRAPeS (red); Trimmed data with TraCeR (gray). b) Same as A, except data was trimmed to include the outer 30bp (instead of 25bp) from each read. Evidently, the successful reconstruction rate of TRAPeS with short reads (average of 84.7% with 30bp reads and 82.6% with 25bp reads) is comparable to that of TraCeR with the original 100bp reads (average of 86.3%), and well above that of TraCeR with the trimmed short reads (0% and 26.7% for 25bp and 30bp respectively).

**Supplementary Figure 5:**
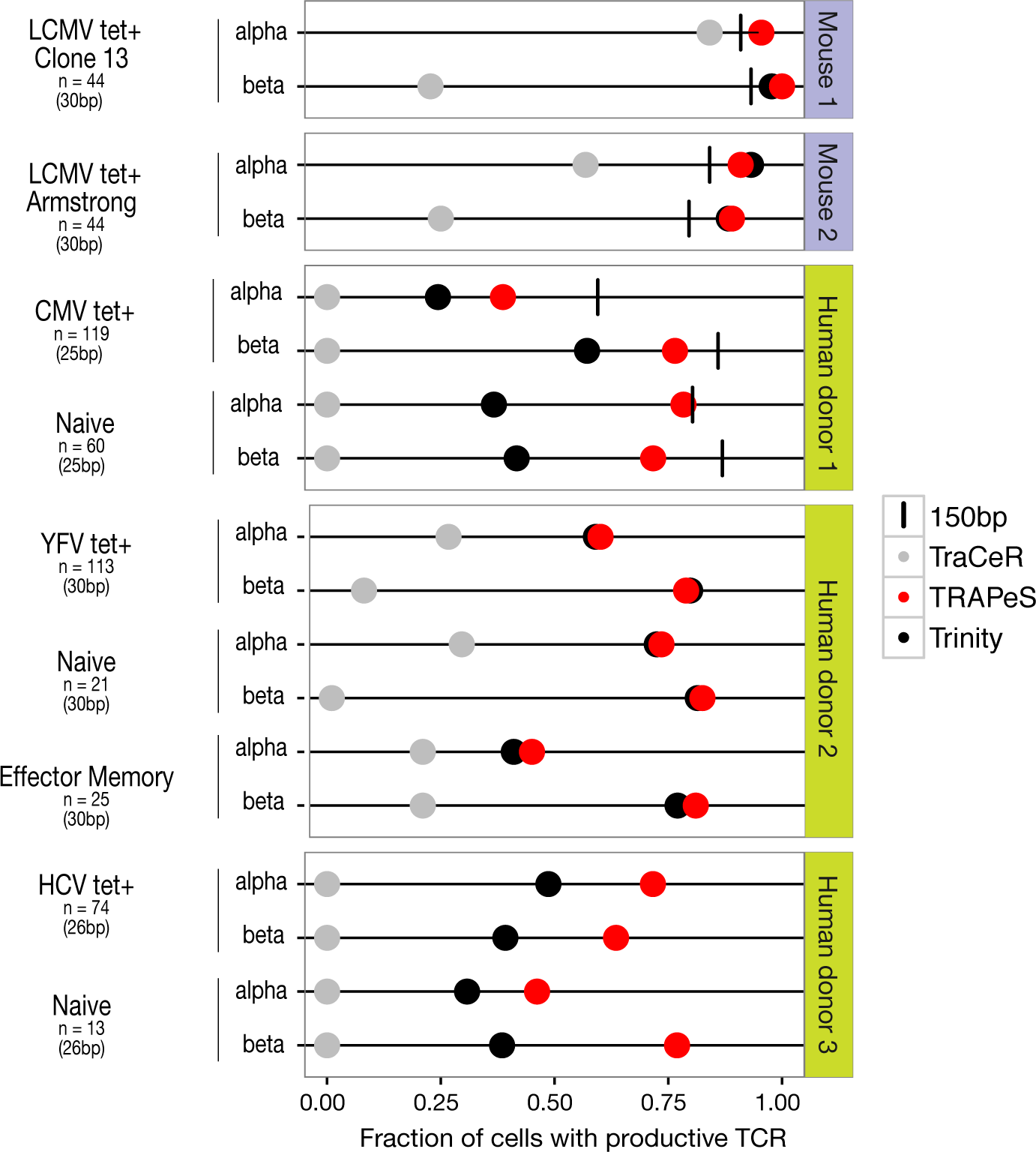
Success rates for reconstruction of productive CDR3 in the various CD8^+^ T cell data sets described in figure 1b, after applying cell-quality filtering as in the TraCeR paper (Stubbington et al). Results are based on the same data as in Figure 1b, with the exception of being limited only to cells that passed the quality criteria used by Stubbington et al. (filtered libraries are those that captured less than 2000 genes, or had more than 10% of reads mapping to mitochondrial genes). Each line depicts the fraction of cells with a productive alpha or beta chain in a given data set with each one of the following methods - 150bp sequencing (black line), short paired-end data reconstructed using TRAPeS (red), TraCeR (light grey) or Trinity (black). Evidently, after cell-quality filtering, the average rate of successful reconstructions in our mouse libraries is 93.7% (with 25-30bp reads), which is higher than that achieved with mouse libraries in the TraCeR paper (86.3%; with 100bp reads [Figure S4 and Stubbington et al.]). No human libraries were analyzed in Stubbington et al.

**Supplementary Figure 6:**
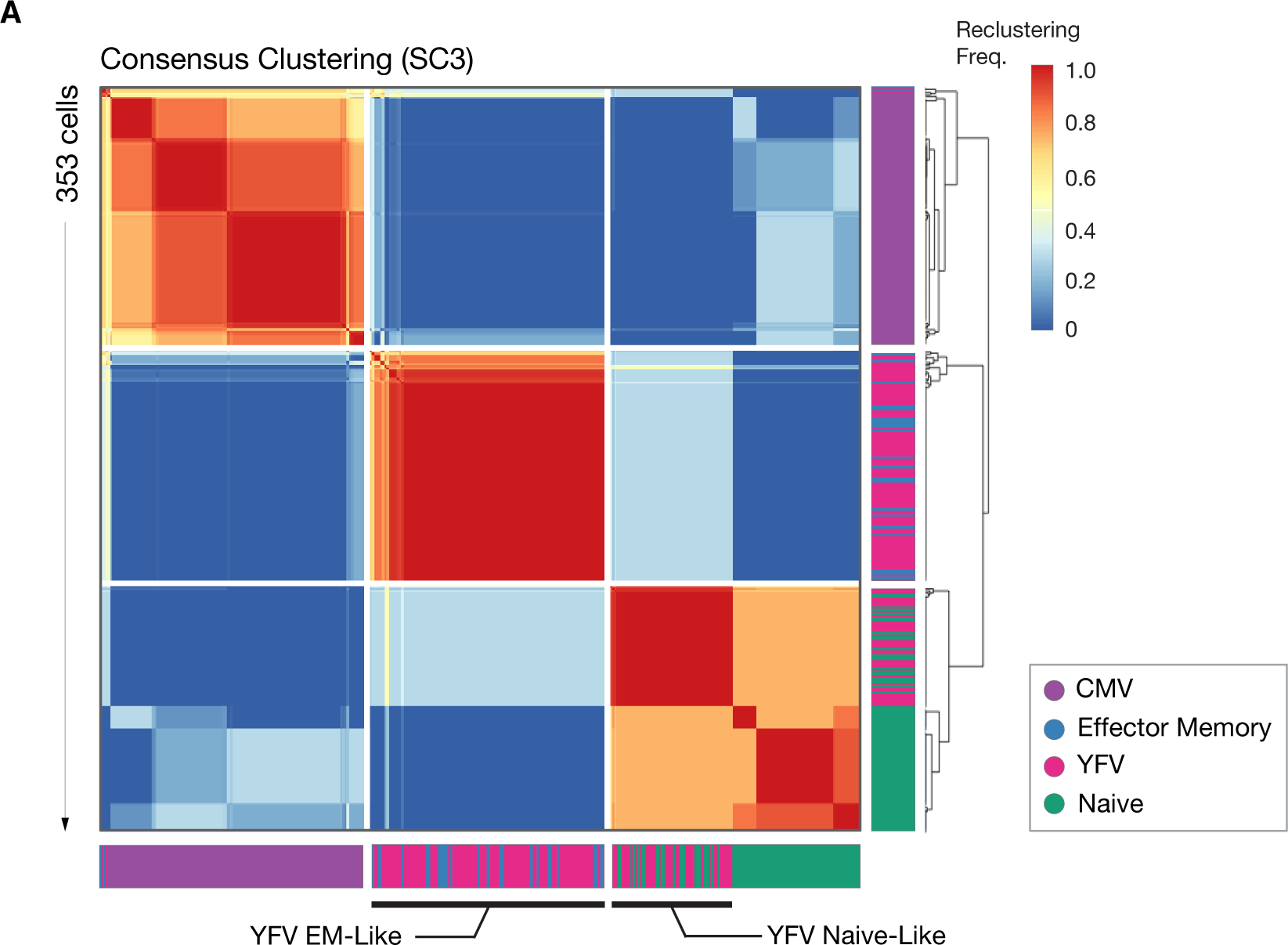
SC3 consensus clustering matrix -. Heatmap of pairwise reclustering frequencies between 353 single cells, indicating three unsupervised clusters. Color bars indicate cellular phenotypes. YFV-specific cells co-clustering with Effector Memory cells were subsequently annotated as Effector Memory-like, and YFV-specific cells co-clustering with true naive cells were annotated as Naive-like.

**Supplementary Figure 7:**
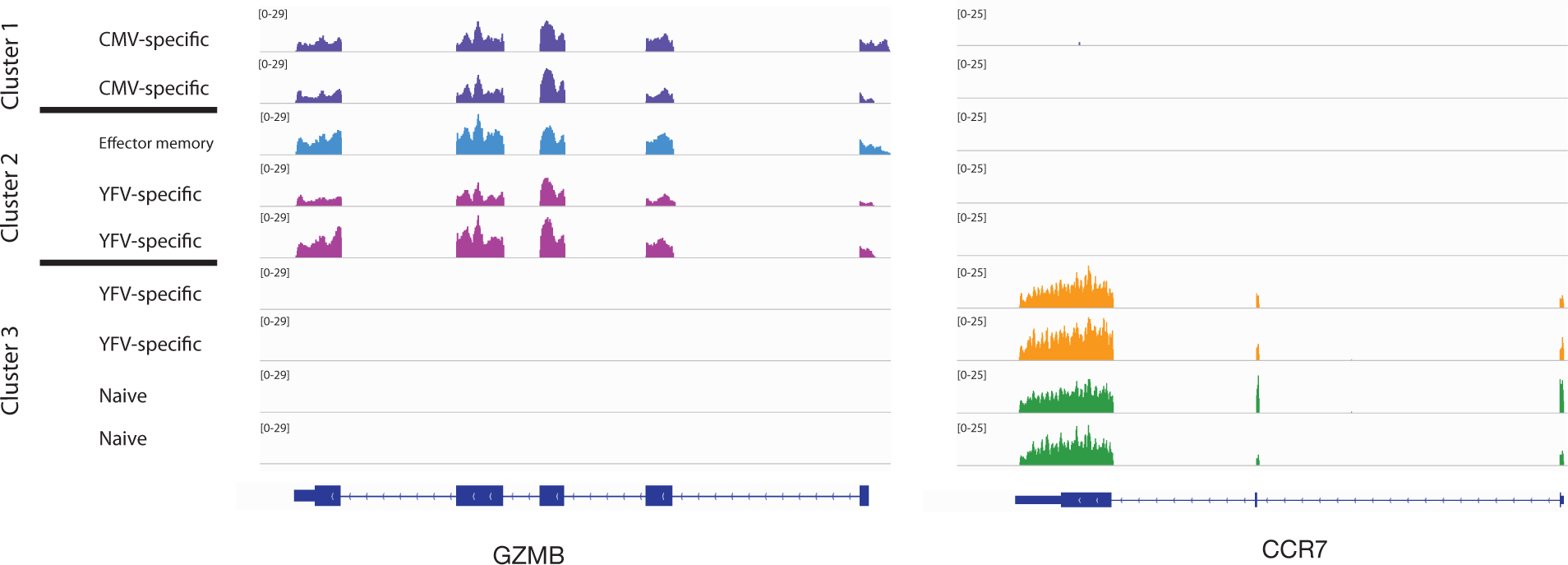
Gene expression changes across clusters -. An Integrative Genome Viewer plot of selected cells from all clusters of genes highly expressed in naive cells (CCR7, right) and antigen-experienced cells (Granzyme B, left).

**Supplementary Figure 8:**
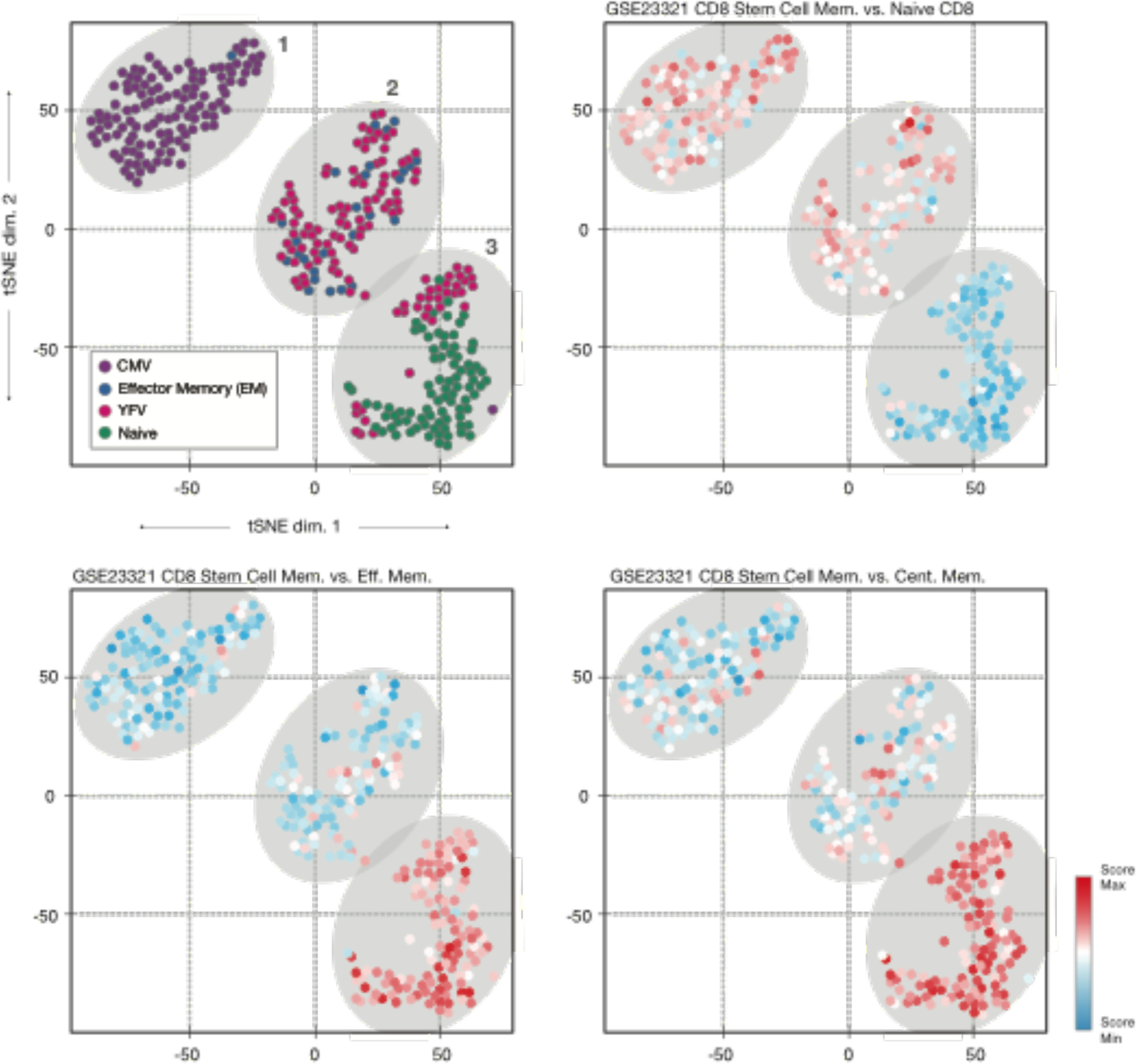
Stem cell signatures -. t-SNE projections, cells colored by relative signature score for a CD8 Stem Cell Memory vs. Naive, CD8 Stem Cell Memory vs. Effector Memory, and CD8 Stem Cell Memory vs. Central Memory signature from ImmuneSigDB.

